# L-Norvaline, a New Therapeutic Agent against Alzheimer’s disease

**DOI:** 10.1101/491480

**Authors:** Baruh Polis, Kolluru D Srikanth, Vyacheslav Gurevich, Hava Gil-Henn, Abraham O. Samson

## Abstract

Alzheimer’s disease (AD) is a slowly progressive neurodegenerative disorder with an insidious onset. The disease is characterized by cognitive impairment and a distinct pathology with neuritic plaques and neurofibrillary tangles.

Growing evidence highlights the role of arginase activity in the manifestation of AD. Upregulation of arginase was shown to contribute to endothelial dysfunction, atherosclerosis, diabetes, and neurodegeneration. Regulation of arginase activity appears to be a promising approach for interfering with the pathogenesis of AD and other metabolic disorders. Therefore, the enzyme represents a novel therapeutic target.

Here, we administer an arginase inhibitor L-norvaline to a mouse model of AD. Then, we evaluate the neuroprotective effect of L-norvaline using immunohistochemistry, proteomics, and quantitative polymerase chain reaction assays. Finally, we identify the biological pathways activated by the treatment.

Remarkably, we find that L-norvaline treatment reverses the cognitive decline in AD mice. We show the treatment is neuroprotective as indicated by reduced beta-amyloidosis, alleviated microgliosis, and TNFα transcription levels. Moreover, elevated levels of neuroplasticity related protein PSD-95 were detected in the hippocampi of mice treated with L-norvaline. Furthermore, we disclose several biological pathways, which are involved in cell survival and neuroplasticity and are activated by the treatment.

Through these modes of action, L-norvaline has the potential to improve the symptoms of AD and even interfere with its pathogenesis. As such, L-norvaline is a promising neuroprotective molecule that might be tailored for the treatment of a range of neurodegenerative disorders.

## Introduction

Alzheimer’s disease (AD) is the primary cause of dementia presenting a severe health problem for the elderly and a growing economic burden for society. AD-associated dementia is characterized morphologically by neuritic plaques containing amyloid-beta (Aβ) peptide and neurofibrillary tangles that are aggregates of hyperphosphorylated tau protein (Schaeffer et al., 2011). The currently dominant amyloid hypothesis of AD pathogenesis postulates that accumulation of Aβ in the brain is the principal cause of AD development (Selkoe and Hardy, 2002). Several therapeutic protocols, predicated upon the hypothesis, have been trialed in AD patients. Unfortunately, all finalized experiments have failed to improve the cognitive functions of AD patients. Moreover, some trials were terminated because of adverse effects (Lahiri et al., 2014; Doody et al., 2013).

The failure to meet the clinical end-points with the traditional anti-amyloid approaches led to rigorous investigations of other AD-related histopathologies, including microgliosis and astrogliosis. These features characterize AD in a region- and time-dependent manner (Osborn et al., 2016). Moreover, activated microglia produce pro-inflammatory mediators in response to Aβ, which in turn activate astrocytes and eventually lead to neuropathology (Heneka et al., 2015). Therefore, dysregulation of cytokine signaling contributes to AD-associated learning and memory deficiency and synaptic dysfunction (Morris et al., 2013). Successful attempts to manipulate the microglial proliferation in order to treat AD-associated memory deficiency in rodents have been achieved recently (Spangenberg et al., 2016).

Contemporary studies indicate that urea cycle disordering plays a role in the pathogenesis of AD (Kan et al., 2015). Specifically, the brains of AD patients show significantly upregulated arginase activity, and reduced nitric oxide synthase (NOS) activity (Liu et al., 2014), which followed by a decrease in the levels of their mutual substrate - L-arginine (Gueli and Taibi, 2013). Moreover, the brains of AD patients contain significantly higher concentrations of urea, which is a waste product of the reaction catalyzed by arginase, compared to healthy brains (Xu et al., 2016a).

Our previous report associated local amyloid-beta-driven immune-mediated response with altered L-arginine metabolism and suggested that arginase inhibition by L-norvaline interrupts the progression of AD (Polis et al., 2018). Accordingly, we hypothesize that dysregulation of the urea cycle in the brain, with upregulation of arginase and consequent nitric oxide (NO) and L-arginine deficiency, leads to the clinical manifestation of AD pathology. Additionally, we suggest that brain overload with waste products (i.e., urea) is a critical leading to neuroinflammation component of the AD pathogenesis. Therefore, arginase is a potential target for treating AD.

To evaluate our hypothesis, we use an arginase inhibitor, L-norvaline, in a murine model of AD and apply a set of immunohistochemistry, proteomics, and quantitative polymerase chain reaction assays. Bioinformatics analysis is utilized to identify the involved biological pathways. We show that animals, treated with L-norvaline, display a significant memory improvement. Additionally, the treatment leads to a decrease in hippocampal Aβ burden, followed by a substantial reduction of microgliosis with a decrease in transcription levels of the TNFα gene. We report elevated levels of neuroplasticity related postsynaptic density protein 95 (PSD-95) in the hippocampi of the animals treated with L-norvaline, which concurs with our earlier reports of increased hippocampal spine density. Finally, we disclose several biological pathways that are involved in cell survival and neuroplasticity that are activated by the treatment. Collectively, out data suggest that L-norvaline could be used as a therapeutic agent in AD.

## Materials and Methods

### Strains of Mice and Treatment

Homozygous triple-transgenic mice (3×Tg) harboring PS1(M146V), APP(Swe), and tau(P301L) transgenes were purchased from Jackson Laboratory (Bar Harbor, ME) and bred in our animal facility. These animals exhibit a synaptic deficiency, plaque, and tangle pathology (Oddo et al., 2003).

Randomly chosen four-month-old homozygous 3×Tg mice were divided into two groups (13 mice in each group). The animals were housed in cages (5 mice per cage) and provided with water and food *ad libitum*. The control animals received regular water. The experimental mice received water with dissolved L-norvaline (Sigma) (250 mg/L). The experiment lasted 2.5 months. The ethics committee for animal experiments of Bar-Ilan University approved the experimental protocol (Permit Number: 82-10–2017).

### Fear Conditioning Test

Contextual fear conditioning is a form of associative learning, in which an animal learns to associate the neutral conditioned stimulus, with the presence of a motivationally significant unconditioned stimulus, such as an electric shock (Wehner and Radcliffe, 2004). In this paradigm, the freezing behavior represents a typical natural response in rodents. The easily assessable lack of movement provides a readout of the memory acquisition and reflects the integrity of the hippocampus (Anagnostaras et al., 2001).

The experiments were performed using two standard chambers with shock floors purchased from Noldus^®^. Each chamber had a 17 × 17 cm floor and is 30 cm in height and was equipped with a top unit including a matrix of infrared lights and an infrared camera, with a high-pass filter blocking visible light. The floor of the chambers included a stainless steel grid (inter-bar separation 0.9 cm) connected to an electric shock generator. Automated tracking was done with EthoVision XT 10 software also provided by Noldus^®^. Mice were handled for three successive days for 5 min a day. During day one, an animal was placed in a chamber for five min and exposed to white background noise. On the second day of a conditioning session, mice received two 2 sec long 0.75 mA foot shocks, at 3.0, and 4.5 min after placement into the chamber. On the third day of the testing session, mice were exposed for five min to the same conditioning context without a shock. EthoVision software controlled the shock periods and amplitude, and the experimental parameters such as trial time and testing zones via user predefined variables.

### Tissue Sampling

The animals were anesthetized with pentobarbital and decapitated. Sections (0.5 mm thick) between 1.7 mm and 2.2 mm posterior to bregma (by the atlas of Franklin and Paxinos) were collected for sampling (Franklin and Paxinos, 2008). The hippocampi were perforated at the dentate gyrus with a 13-gauge dissection needle. The brain punched tissues were frozen, and stored at minus 80° C.

### Pathway Enrichment Analysis

In order to identify specific biomarkers and signatures of phenotypic state caused by L-norvaline treatments in 3×Tg mice as a biological system, we accomplished a functional interpretation of the genes derived from the antibody microarray assay done in the previous study (Polis et al., 2018). The array includes 1448 targets. There are 84 significantly up and down-regulated proteins with significant (p-value<0.05) change of expression in the rate more than 25% (Supplementary Table S1). The pathway enrichment analysis was performed using Ingenuity^®^ Pathway Analysis software (IPA^®^).

### Immunostaining

Four animals from each group were deeply anesthetized and transcardially perfused with 30 ml of phosphate buffer saline (PBS), followed by 50 ml of chilled paraformaldehyde 4% in PBS. The brains, liver, and kidney were carefully removed and fixed in 4% paraformaldehyde for 24 h. The organs were transferred to 70% ethanol at 4° C for 48 h, dehydrated and paraffin embedded. The paraffin-embedded blocks were ice-cooled and sliced at a thickness of 4 μm. The sections were mounted, dried overnight at room temperature, and stored at 4° C.

For the quantitative histochemical analysis of beta-amyloid, three coronal brain sections cut at 25 μm intervals throughout the brain per mouse (1.8–2.0 mm posterior to bregma) were used. Immunohistochemistry (IHC) was performed on the plane-matched coronal sections.

The kidneys were sliced in half longitudinally, and the liver was cut in transverse sections. For the analysis of arginase 1 (ARG1) immunopositivity in the liver and arginase 2 (ARG2) in the kidney, two 4 μm sections were cut per organ at 100 μm intervals.

Staining was accomplished on a Leica Bond III system (Leica Biosystems Newcastle Ltd., UK). The pretreated with an epitope-retrieval solution (Leica Biosystems Newcastle Ltd, UK) tissues were incubated with primary antibodies for 30 min. For the 6E10 (Abcam, #ab2539), arginase 1 (ARG1) (GeneTex, #GTX113131), arginase2 (ARG2) (GeneTex, #GTX104036) antibodies dilutions were 1:200, 1:500, 1:600 respectively. A Leica Refine-HRP kit served for hematoxylin counterstaining. The omission of the primary antibodies served as a negative control.

### Imaging and Quantification Analysis

The sections were viewed under a slide scanner Axio Scan.Z1 (Zeiss, Oberkochen, Germany) with a 40×/0.95 objective. The images were captured at various focal distances (Z-planes) to provide a composite image with a greater depth of field (mostly at 0.5 μm). The image analysis was carried out using Zen Blue 2.5 (Zeiss, Oberkochen, Germany). A fixed background intensity threshold was set for all sections representing a single type of staining. For 6E10 staining, the analysis of immunopositivity was performed in the CA1, CA3, CA4 areas of the hippocampus. The surface of the immunopositive area (above the threshold) has been subjected to the statistical analysis.

Additionally, the image densitometry method was applied to quantify the amount of staining in the specimens. Integrated Optical Density (ODi), which is the optical density of individual pixels in the image reflecting the expression levels of the proteins, was measured via digital image analysis software (Image-Pro^®^ 10.0.1; Media Cybernetics, Inc., Rockville, MD), and presented as the average value for each treatment group.

The glomerular area was quantified using ZEN Blue 2.5. Twenty-four cortical glomeruli were measured in each treatment group. In order to ensure that chosen glomeruli were sectioned in the same plane, only those containing evident efferent and afferent arteriolar stalks have been analyzed. Bowman’s capsule diameters were statistically compared.

### RNA Extraction, Reverse Transcription, and Real-time Polymerase Chain Reaction

Hippocampal tissues were sampled from five animals (each group). Total RNA was isolated using the RNeasy Mini Kit (Cat. No. 74104, Qiagen) following the manufacturer’s instructions including DNase treatment. RNA quantification was performed using Qubit™ RNA HS Assay Kit (Cat. No. Q32852, Invitrogen). RNA integrity (RIN) was measured using Agilent 2100 Bioanalyzer System and Agilent RNA 6000 Pico Kit (Cat. No. 5067-1513, Agilent Technologies). cDNA was prepared from 200ng of total RNA using SuperScript^®^ III First-Strand Synthesis System for real-time polymerase chain reaction (RT-PCR) (Cat. No. 18080-051, Invitrogen) following the manufacturer’s instructions. Real-time PCR was performed using TaqMan probes (Applied Biosystems). TNF RNA levels were analyzed with Tnf: Mm00443258_m1 probe. For the normalization of TNF RNA levels, ACTB and GAPDH endogenous housekeeping genes controls were analyzed using ActB: Mm00607939_s1 and Gapdh: Mm99999915_g1 probes respectively. PCR was set in triplicates following the manufacturer’s instructions (Applied Biosystems, Insert PN 4444602 Rev. C) in a ten μl volume using five ng cDNA template. PCR was run, and the data was analyzed in the StepOnePlus system installed with StepOne Software v2.3. The quantification was performed using the comparative Ct (ΔΔCt) method.

### Western Blotting

Hippocampal tissue was homogenized in an equal ratio of tissue homogenate buffer (50 mM Tris-HCl (pH 7.5), 150 mM KCl, 0.32 M sucrose, Protease inhibitor cocktail (Sigma)) and lysis buffer (1%Triton, 10% glycerol, 120mM NaCl, 25mM HEPES, 1mM EDTA, 0.75mM MgCl2, 2mM NaF, 1mM Sodium vanadate & Protease inhibitor). The samples were incubated on ice for 10 min and centrifuged at 11,000g for 10 minutes at 4°C. The supernatant was aliquoted, and protein concentration was determined using a protein assay kit (BioRad). Forty μg samples were subjected to SDS-PAGE and transferred onto a nitrocellulose membrane. The membranes were blocked for one hour at room temperature in TBS containing 0.1% casein. The membranes were incubated overnight with primary antibody 1:500 dilution of PSD-95 (Abcam, ab18258), BACE-1(Abcam, ab2077), APP (Abcam, ab2072). Following washing with TBST, the membranes were incubated with LI-COR dye-conjugated secondary antibody for one h. The beta-actin antibody was used as a reference (Abcam, ab8227) at 1:5000 dilution. Membranes were scanned on the LI-COR Odyssey scanner.

### Statistical Analysis

Statistical analyses were conducted with GraphPad Prism 7.0 for Windows (GraphPad Software, Inc.). The significance was set at 95% of confidence. The two-tailed Student’s *t*-test was performed to compare the means. All data are presented as mean values. Throughout the text and in plots, the variability is indicated by the standard error of the mean (SEM).

## Results

### L-Norvaline Improved Memory Acquisition in the 3×Tg Mice

Contextual fear conditioning paradigm was utilized to study the effect of L-norvaline treatment upon the rate of learning and memory functions in the 3xTg mice. Initial habituation period (Fig. 1A) on the first day allowed acclimation to the experimental environment and reduction of orienting responses. Freezing behavior of the mice was tested 24 hours after training. The animals treated with L-norvaline exhibited significantly increased freezing time during the fourth (66.7±4.96% vs. 51.8±4.25%, p-value=0.033) and fifth (63.84±5.8% vs. 45.4±4.3%, p-value=0.018) minutes of the test compared to the control group (Fig. 1C). Taking into account that the experimental animals were exposed to the shocks only during the second part of the procedure, we conclude that their contextual memory acquisition, which is associated with the conditioned stimulus, improved significantly following the treatment with L-norvaline.

**Figure 1.**
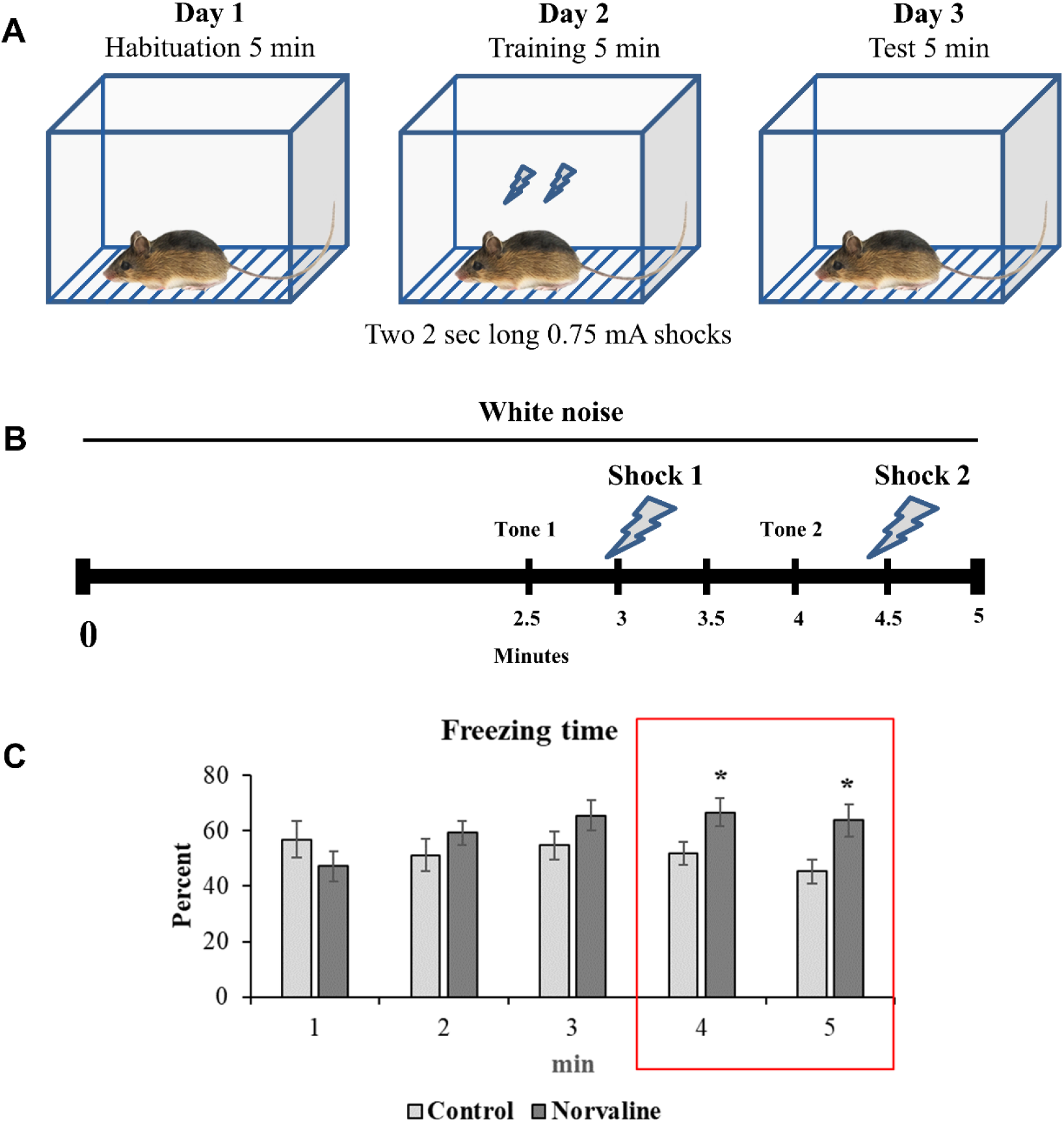
Contextual fear conditioning. (A) Experimental design (B) Fear training scheme. (C) Effect of L-norvaline treatment on freezing time during the five-minute test. Mean freezing levels (± SEM) of controls (n = 13), and L-norvaline treated mice (n = 13). The two-tailed Student’s *t*-test revealed a significant effect of the treatment on freezing time, *p<0.05.

### L-norvaline Reduced β-Amyloidosis in the Hippocampi of 3×Tg Mice

Coronal brain sections of the 3×Tg mice were stained with anti- APP/Aβ-specific antibody to study the impact of the L-norvaline treatment on the amyloid burden. Six- to seven-month-old 3×Tg mice exhibit enhanced intracellular and extracellular deposition of Aβ in their hippocampi (Fig. 2A, B)). Intracellular deposits are most prominent in the CA1 area (Fig.2C), with a substantial reduction in the L-norvaline treated group (Fig.2D). There were only a few plaques detected within the hippocampi. The analysis of immunopositive surface area was performed in the CA1 (Fig.2C-D), CA3 (Fig.2E-F), and CA4 regions of the hippocampi. We detected significant differences in the levels of immunopositivity between the two experimental groups. There was a significant reduction in the 6E10 positivity in the CA1 (p=0.0095), CA3 (p=0.016), CA4 (p=0.016) areas. These data point to an effect of L-norvaline on the level of β-amyloidosis in the 3×Tg mice and support our previously reported findings (Polis et al., 2018).

**Figure 2.**
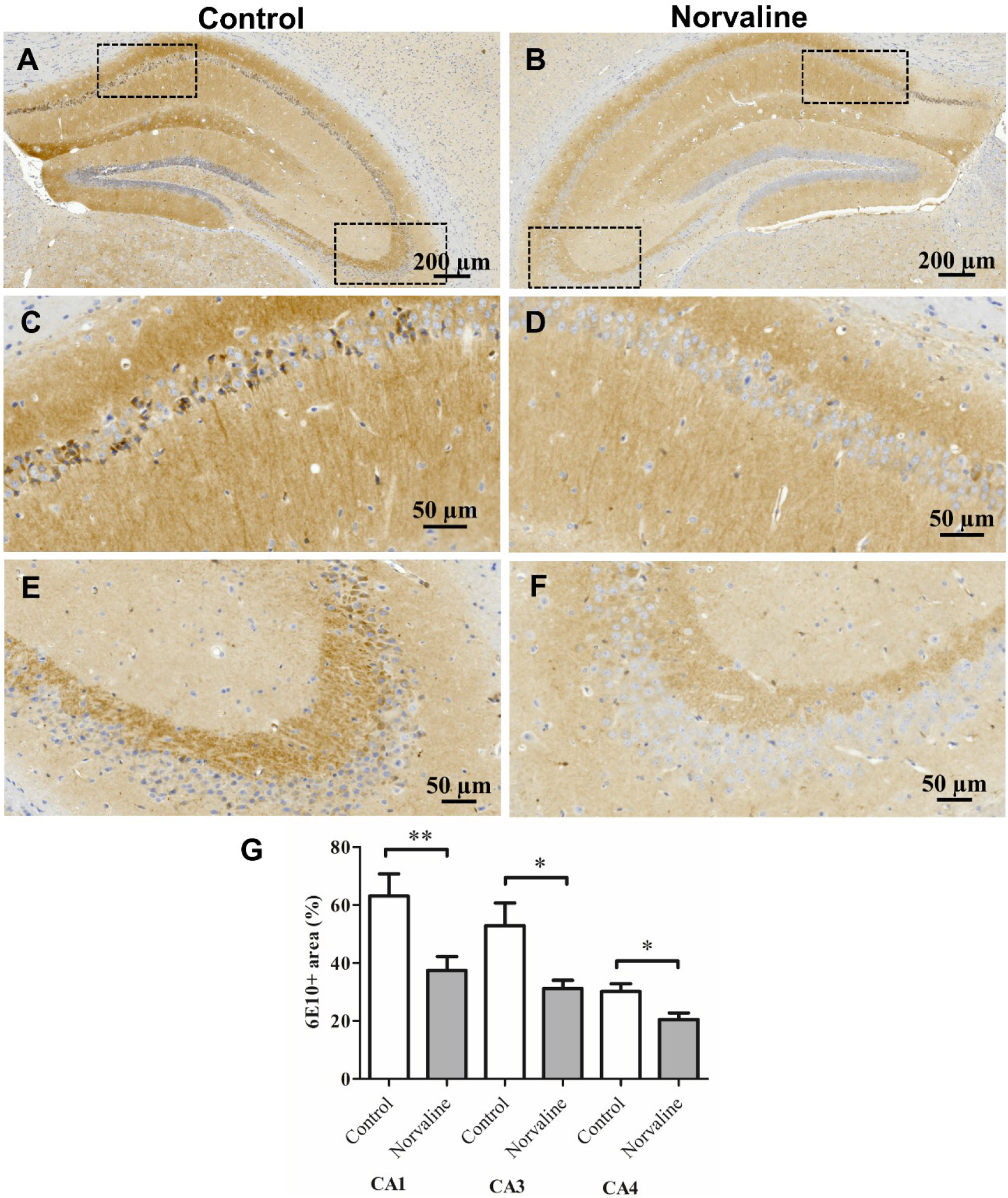
The effect of L-norvaline upon the hippocampal Aβ burden in the 3×Tg mice. Representative ×40 photomicrographs of the entire hippocampi stained with 6E10 antibody from the control (A) and L-norvaline treated (B) mice. Insets (C) and (D) represent the CA1 region, (E) and (F) – CA3 region. Quantification of the Aβ immunopositive area in different brain regions of the 3×Tg mice treated with vehicle (control) or L-norvaline (G). The Student t-test was used to compare the means between 2 groups, *p<0.05, **p<0.01 (n = 12, three mice per group).

### L-Norvaline Reduced the Transcription Levels of the TNFα gene

Tumor necrosis factor signaling is required for Aβ-induced neuronal death (Sumbria et al., 2017). The critical role of TNFα in AD pathogenesis is supported by observations made in the mouse models of AD (Sly et al., 2001). Elevated TNFα levels were observed in the brain tissues of AD transgenic mice. In order to investigate the impact of L-norvaline treatment on TNFα gene expression levels, mRNA was isolated from the hippocampi of control and treated mice, and analyzed by quantitative RT-PCR. cDNA levels were normalized to GAPDH, and AKT as reference genes and presented relative to gene expression in vehicle controls. The two-tailed Student’s *t*-test (t=2.66, df=8) revealed a significant (p=0.029) reduction (by 35.5%) of TNFα gene expression levels in the L-norvaline treated group (Fig. 3).

**Figure 3.**
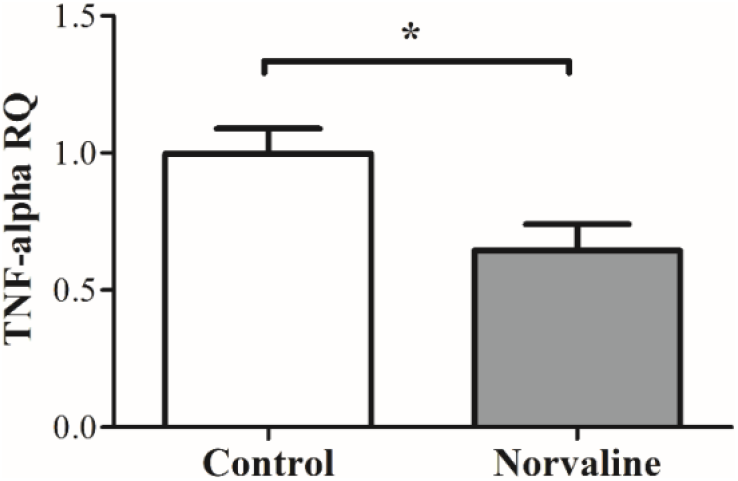
TNFα mRNA expression levels. Real-time PCR analysis of mRNA levels of TNFα gene. All data are presented as the mean and SEM (n = 5). *p<0.05; (*t*-test).

### L-Norvaline Amplified the Expression Levels of Postsynaptic Density-95 Protein in the 3×Tg Mice

The impact of the L-norvaline treatment upon the hippocampal levels of neuroplasticity-associated protein PSD-95 in the 3×Tg mice was investigated using western blotting. Decreased levels of PSD-95 in the cerebral cortex and hippocampus of 3×Tg mice compared to the wild-type (WT) mice was previously reported (Revilla et al., 2014a) suggesting synaptic integrity loss in this model. We have analyzed previously the effect of L-norvaline upon the spine density and the expression levels a list of neuroplasticity related proteins and disclosed a relative spine deficiency in the 3×Tg mice compared to WT and evidenced an increase in spine density (by about 20%) following the treatment (Polis et al., 2018).

In order to evaluate further the effect of the treatment on synaptic integrity, we measured the levels of the post-synaptic protein PSD-95. We observed a significant (p-value=0.015) increase (by 17.2%) in PSD-95 levels in the hippocampi of 3×Tg mice treated with L-norvaline compared to the control animals (Fig. 4A, B), which accords with the data acquired by Golgi staining analysis and reported previously.

**Figure 4.**
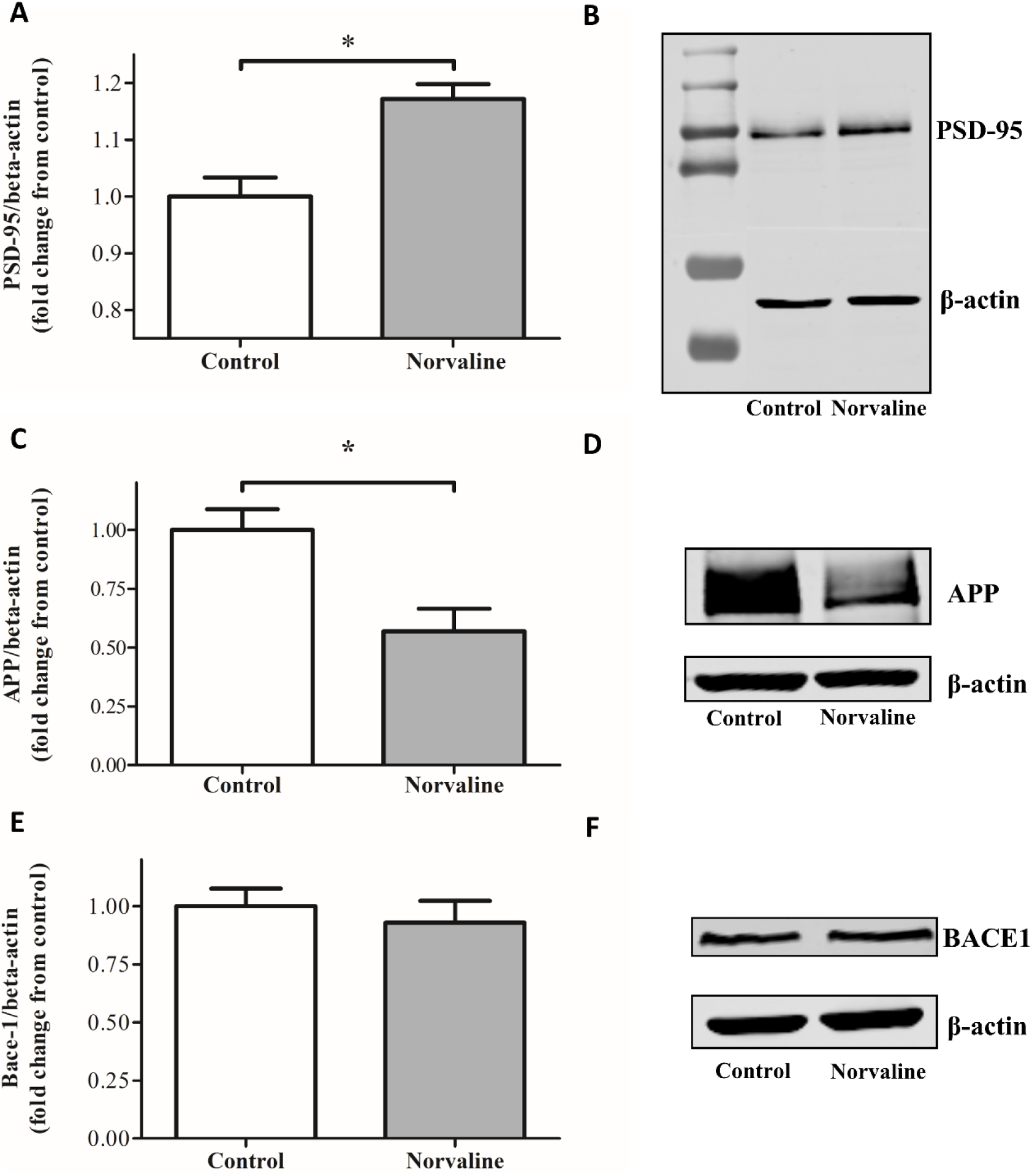
Western blot analysis of the hippocampal lysates using anti-PSD-95, anti-APP, anti-BACE-1, and anti-β-actin antibodies. Immunoreactive band intensities normalized with their respective β-actin bands were compared and presented by bar charts as fold change from vehicle-treated controls. PSD-95 expression levels are significantly higher and APP lower in the brains of L-norvaline mice. There is no significant effect of the treatment upon the levels of BACE-1 protein (data are mean±SEM, *p < 0.05, *t*-test, n=3).

### L-Norvaline Reduced the Expression Levels of APP but not BACE-1 in the 3×Tg Mice

The brains of 3×Tg mice contain growing with age APP levels, which are higher than in WT animals (Croft et al., 2017). Immunohistochemistry with 6E10 antibody revealed a significant reduction of the total amyloid burden in the cortices and hippocampi of mice treated with L-norvaline. In order to comprehend the phenomenon and address it to the increased transcription of APP or altered processing by BACE-1, we performed western blot analysis of the extracted proteins. Quantitative analysis of the blots demonstrated that L-norvaline significantly (p-value=0.029) reduced (by 43%) the levels of APP expression (Fig. 4C, D), however, it had little or no effect (p-value=0.59) on the levels of BACE-1 protein (Fig. 4E, F).

### L-Norvaline Induces Cellular Pathways Involved in Neuroplasticity and Oxidative Stress Protection

Functional interpretation of the genes derived from the antibody microarray assay was performed using Ingenuity^®^ Pathway Analysis software (IPA^®^). The IPA^^®^^ tool was applied to uncover the significance of proteomics data and identify candidate biomarkers within the context of 3×Tg mice as a biological system. The array includes 1448 targets. There were 84 significantly up and down-regulated proteins with the change of expression by more than 25% detected by the assay (Supplementary Table S1).

The analysis with a preset threshold of 25% change from control and the significance set at 95% of confidence revealed that treatment of 3×Tg mice with L-Norvaline led to activation of 41 critical biological processes (Fig. 5). Among the most significant pathways (with p-value<0.0025 and overlap>30%) detected by IPA^®^ were neuregulin pathway, synaptic long-term depression, ERK/MAPK pathway, and oncostatin M pathway. Moreover, IPA^®^ points out several signaling pathways with significant effect, which are involved in cell survival and immune response. Among them are PI3K/AKT signaling, TREM1 signaling, CDK5 signaling, IL-2, IL-3, IL-15 signaling.

**Figure 5.**
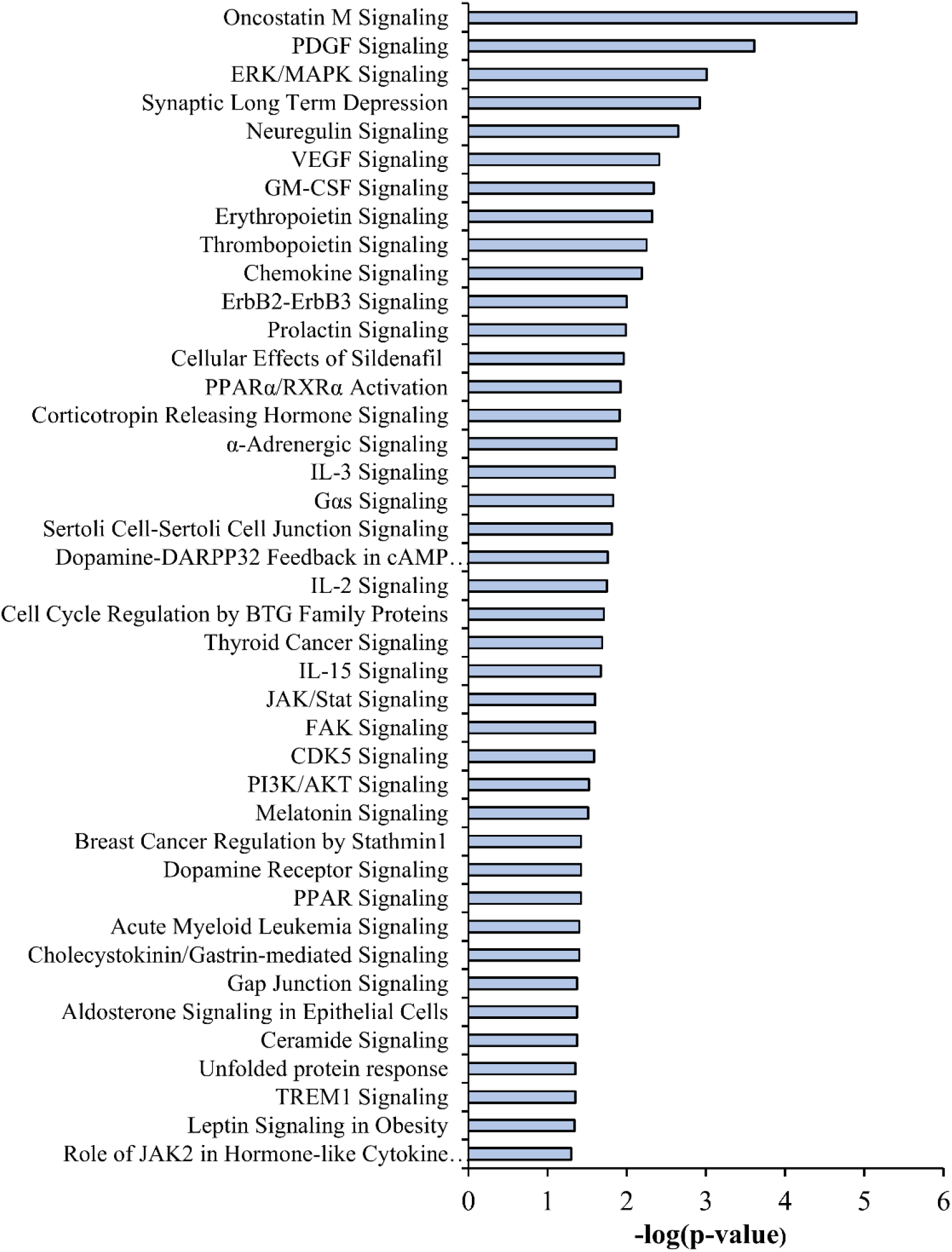
A list of the biological pathways induced by L-norvaline treatment and identified by IPA^®^.

Neuregulins comprise a cluster of epidermal growth factor-like proteins that act on the epidermal growth factor receptor (EGFR) family of receptors and are highly implicated in neural development and brain homeostasis (Mei and Nave, 2014). Accumulating evidence suggests that NRG1 signaling has an impact on cognitive function and neuropathology in AD. Overexpression of NRG1 in the hippocampus of AD mice improves memory deficits and ameliorates disease-associated neuropathology (Xu et al., 2016b). Our results reveal upregulation of several important downstream to NRG proteins. We demonstrate that the levels of Src proto-oncogene-encoded protein-tyrosine kinase increase significantly (p-value=0.003) by 94% in the brains of the treated group. Src activity was shown to be required for avoidance memory formation and recall (Bevilaqua et al., 2003). Moreover, the levels of Signal Transducer and Activator of Transcription 5A (STAT5) escalated significantly (p=0.006) by 45%. STAT5 is an essential cellular mechanism regulating cognitive functions. Brain-specific STAT5 ablation was shown recently to impair learning and memory formation in mice (Furigo et al., 2018). Another study suggests a contribution of STAT5 in neuroprotection (Zhang et al., 2007). The authors demonstrate a strong association between STAT5 activation and neuronal survival in the rat hippocampus after cerebral ischemia.

The platelet-derived growth factor (PDGF) ligands and receptors are expressed during embryonic development and in the mature nervous system. In pathologic conditions, PDGF modulates neuronal excitability and stimulates survival signals, via the PI3-K/Akt pathway and other ways, rescuing cells from apoptosis (Funa and Sasahara, 2014). PDGF signaling defects have been shown to underlie the clinical progression of neurodegeneration in multiple sclerosis (Mori et al., 2013). Our assay confirms that PDGF signaling is highly involved and activated by L-norvaline treatment (p=0.00025). The levels of one of the central enzymes of the pathway, sphingosine kinase 2 (SPHK2), increased significantly (p=0.036) by 56% following the treatment. SPHK2 is highly implicated in AD pathogenesis as indicated by dysregulation of the sphingolipid metabolism (i.e., reduced levels of sphingosine 1-phosphate (S1P), a product of the reaction) in AD brains (Dominguez et al., 2018). S1P is a potent neuroprotective signaling lipid (Couttas et al., 2014), which acts via specific G protein-coupled receptors (S1PRs) and regulates microglial number and activity in the brain (Hla and Brinkmann, 2011).

Vascular endothelial growth factor (VEGF) is essential for neuroprotection (Rosenstein et al., 2010). VEGF signaling activation is particularly beneficial in individuals showing early hallmarks of AD (Hohman et al., 2015). We demonstrate that L-norvaline significantly (p-value=0.004) up-regulates VEGF signaling pathway in the 3×Tg mice. L-norvaline treatment additionally leads to up-regulation of the ERK/MAPK signaling pathway, which is essential for endogenous neuroprotection (Karmarkar et al., 2011). Moreover, IPA^®^ points out other pathways with significant effect, which are involved in cell survival and immune response.

Noteworthy, L-norvaline treatment led to a significant (p-value=0.04) increase (by 53%) in the levels of Glial Cell-Derived Neurotrophic Factor (GDNF) receptor Ret, as detected by the antibody array. Ret is a tyrosine kinase and common signaling receptor for GDNF-family ligands (Airaksinen and Saarma, 2002). Generally, GDNF is an effective supporter of neuronal survival (Allen et al., 2013), and Ret is essential for mediating GDNF neuroprotective and neuroregenerative effects (Drinkut et al., 2016). Moreover, Neural Cell Adhesion Molecule (NCAM), which has been identified as a second signaling receptor for GDNF (Paratcha et al., 2003), demonstrated a significantly (p-value=0.003) elevated (by 43%) levels following the treatment. NCAM regulates synaptic plasticity (Lüthi et al., 1994), and mediates the axonal growth in the hippocampal and cortical neurons (Paratcha et al., 2003). Of note, GDNF is down-regulated in seven-month-old 3×Tg mice (Revilla et al., 2014a) and GDNF overexpression improves learning and memory in this model of AD (Revilla et al., 2014b).

Additionally, L-norvaline treatment led to a substantial increase (by 59%) in the levels of protein-serine kinase suppressor of Ras 1 (Ksr1), which is critical for neuroprotection. Ksr1 mediates the anti-apoptotic effects of Brain-Derived Neurotrophic Factor (BDNF) (Szatmari et al., 2007), which is highly implicated in AD (Peng et al., 2005).

Contemporary data demonstrate that another essential neuroprotective factor, Nerve Growth Factor (NGF), plays a crucial role in normal aging (Parikh et al., 2013) and AD pathogenesis (Iulita and Cuello, 2014). Consequently, NGF application has emerged as a novel approach for AD therapy (Eyjolfsdottir et al., 2016). Of note, the neuroprotective effects of NGF are mediated via Tropomyosin receptor kinase A (TrkA) (Nguyen et al., 2010), which was significantly (p-value=0.039) increased (by 56%) following the treatment.

Antibody microarray detected several other essential for cell survival proteins that demonstrated a significant but moderate (less than 25% cutoff for IPA^®^) escalation rate following the treatment. For example, superoxide dismutase [Cu-Zn] (SOD) levels were elevated by 19% in the brains of the treated animals. This enzyme plays a critical role in cellular response to oxygen-containing compounds and has been shown to be neuroprotective in a rat model with NMDA excitotoxic lesion (Peluffo et al., 2006). Additionally, we observed a 24% increase in the levels of phosphatidylinositol 3-kinase regulatory subunit alpha, which protects against H_2_O_2_-induced neuron degeneration (Yu et al., 2004). Accordingly, we suggest that L-norvaline possesses several converging on neuroprotection modes of activity (Fig. 8).

### L-Norvaline Effectively Reduced the Expression Levels of ARG1 and ARG2 in the Primary Organs of their Activity without Inducing Morphological Aberrations

In order to evaluate the rate of ARG1 inhibition by L-norvaline in the hepatic tissue, we performed a histological analysis of the 3×Tg mice liver. We did not detect structural changes of the classical hexagonal, divided into concentric parts, hepatic lobular structure following the treatment (Fig. 6A-D). However, the levels of ARG1 immunopositivity were significantly reduced in the L-norvaline treated group, which was reflected by a decrease in ARG1 positive surface area (p-value=0.015) and integrated optical density (p-value=0.04).

**Figure 6.**
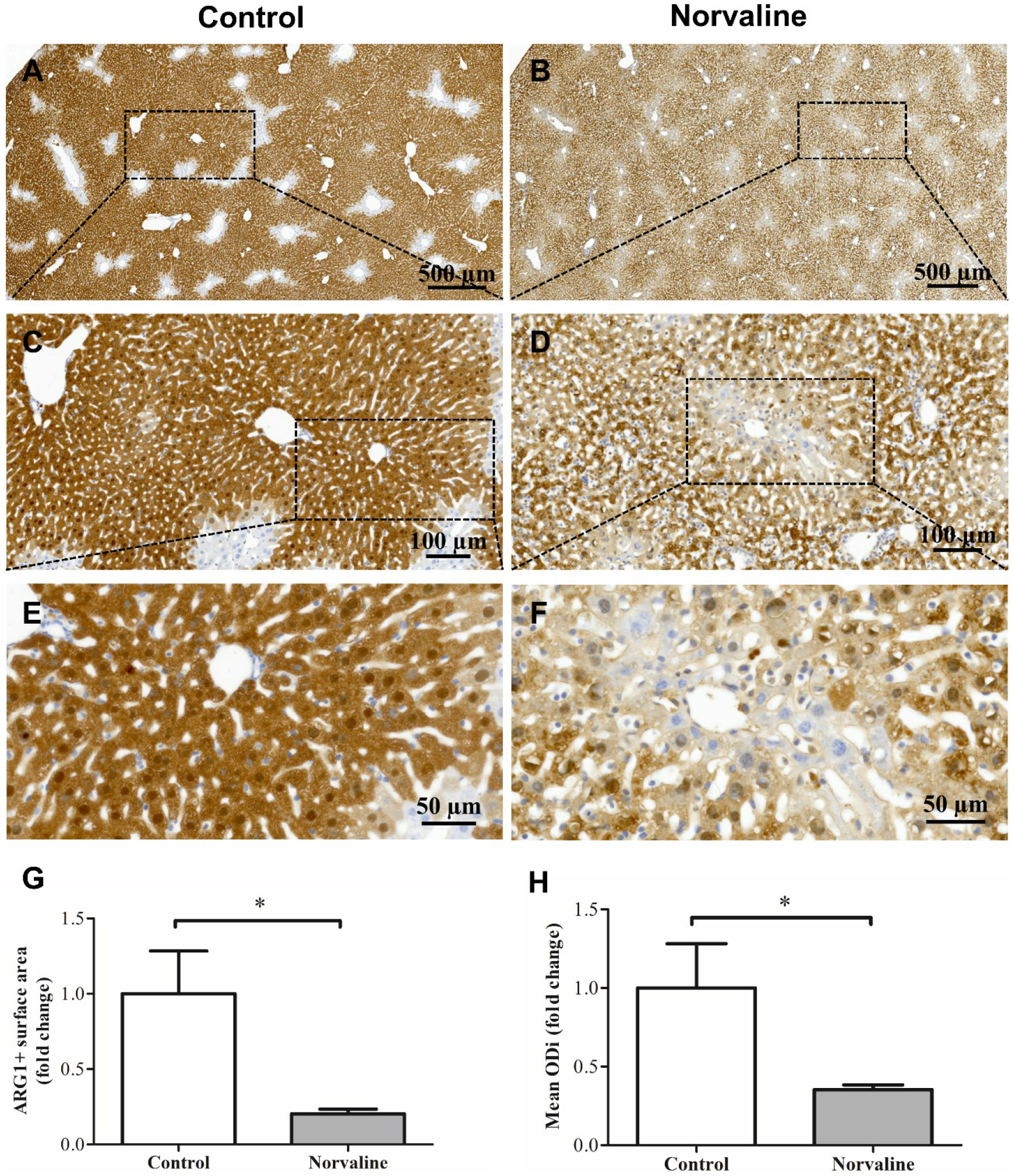
Immunohistochemical ARG1 staining of the 3×Tg mice liver tissue. Bright-field ×40 micrographs of the liver from control (A, C, E) and L-Norvaline treated animals (B, D, F). The bar charts show a significant reduction in the ARG1+ surface area (G) and integrated optical density (ODi) of ARG1+objects in the treated with L-norvaline group (H) (n = 8, four mice per group). Student’s *t-*test *p<0.05

Additionally, we performed renal tissue staining with ARG2 antibody and hematoxylin to assess the kidney structural integrity and the levels of ARG2 expression following the L-norvaline treatment. It was shown previously in the rat, that ARG2 is intensely expressed in the proximal straight tubules of outer medulla, and a subpopulation of the proximal tubules of the renal cortex (Miyanaka et al., 1998). Our data accord with these findings (Fig. 7).

**Figure 7.**
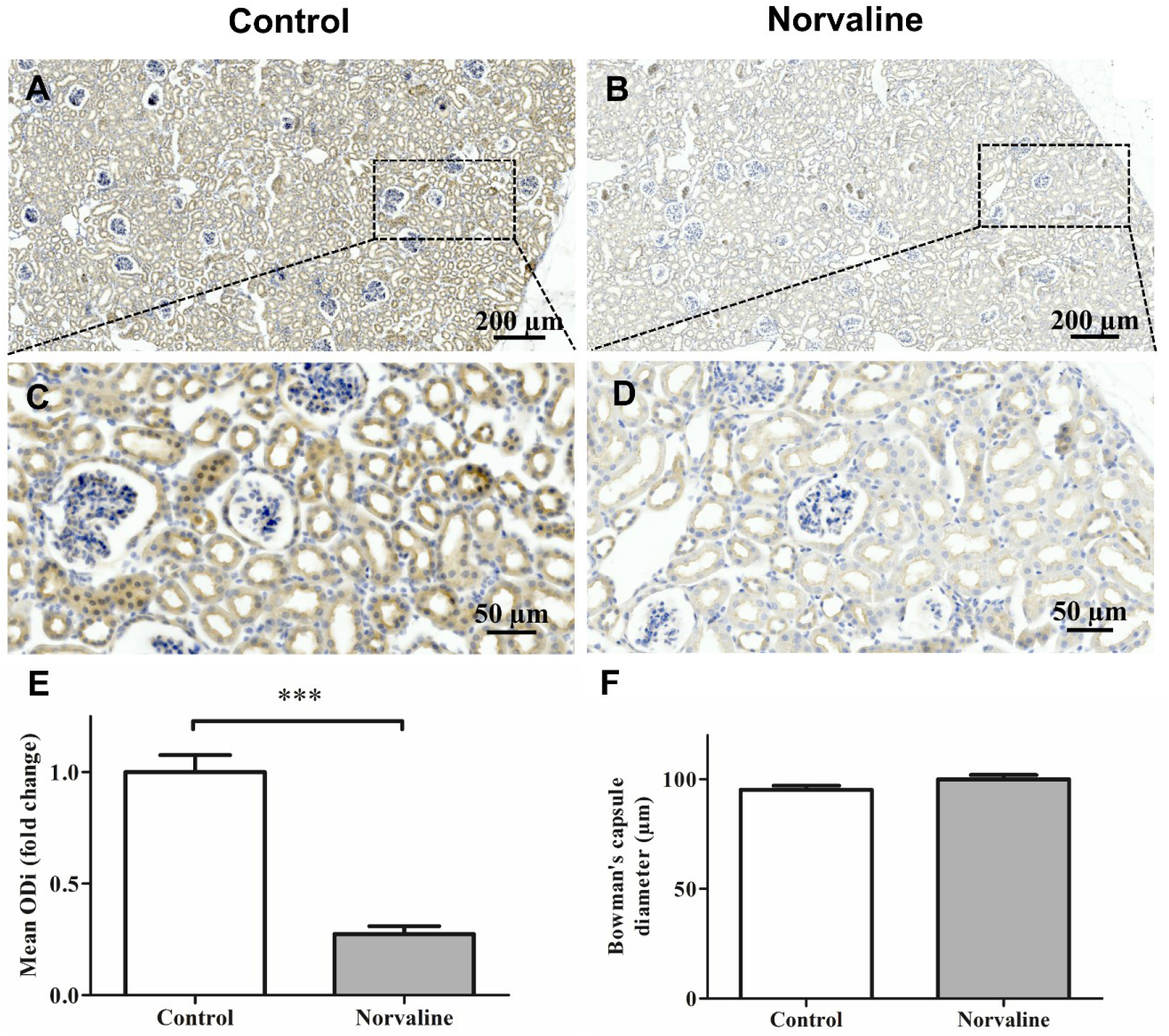
Immunohistochemical ARG2 staining of the 3×Tg mice kidney tissue. Representative ×20 bright-field micrographs of the longitudinally sliced kidneys from the control (A) and L-norvaline treated (B) mice. Insets (C, D) are ×40 images of outer medulla and cortex. The bar charts show a significant reduction in the integrated optical density (ODi) of ARG2-positive objects in the treated with L-norvaline group (E) (n = 8, four mice per group). There was no significant effect on the Bowman’s capsule diameter (F). Student’s *t*-test (n = 24, three cortical objects per section, four mice per group), ***p<0.001

In previous studies, the Bowman’s capsule space expansion was marked as a central histological kidney’s dysfunction characteristic (Thakur et al., 2006; Tobar et al., 2013).

We subjected the cortical glomerular capsule diameter to the statistical analysis to detect deviations in glomerular space. We did not observe any significant variability (p-value= 0.081) in the Bowman’s capsule diameters between experimental groups (Fig. 7F). The mean diameters were 95.15 ± 1.88 μm in the control group and 99.97 ± 1.93 μm in the treated group. Nevertheless, the ARG2 positivity was significantly (p<0.001) reduced in the treated with L-norvaline mice (Fig. 7E).

## Discussion

In this study, we continued the investigation of neuroprotective properties of a non-proteinogenic unbranched-chain amino acid L-norvaline in a mouse model of AD. We proved previously, by use of two canonical paradigms, that L-norvaline ameliorated short- and long-term memory deficits in the 3×Tg mice, and demonstrated a significant reduction of Aβ positivity in the cortices following the treatment (Polis et al., 2018). Here, we confirm the findings with another context-related memory test and report a significant reduction of the hippocampal Aβ burden.

In order to decipher the amyloidosis reduction mechanism, we performed western blotting of hippocampal lysates and studied the effects of L-norvaline treatment upon APP and BACE-1 expression. In the previous report, we revealed a substantial decrease in the levels of Aβ toxic oligomeric and fibrillary forms following the treatment. Here, we evidence a significant reduction of APP, but not BACE-1 protein levels. Therefore, we suggest that L-norvaline moderates the rate of APP translation, which leads to a dramatic decline in the hippocampal amounts of Aβ deposits and toxic oligomers.

Levels of soluble conformations of Aβ have been shown to correlate with AD-associated synaptic loss and cognitive impairment (McLean et al., 1999). Moreover, these species trigger neurotoxicity in the presence of microglia and potentiate proinflammatory changes in the AD brain (Dhawan et al., 2012). Current literature highlights chronic neuroinflammation as a prominent feature of AD pathogenesis (Heneka et al., 2015). Accordingly, there were attempts to interfere with inflammatory complement-mediated processes leading to the synaptic loss in AD mice.

The 3×Tg mice exhibit a regional and age-dependent enhancement of F4/80-positive (Janelsins et al., 2005), IBA1-positive (Montgomery et al., 2011), and CD11b-positive microglia (Ye et al., 2016). The phenomenon is associated with amplification of various factors of inflammation in the brain, including TNFα (Montacute et al., 2017). Microglia and neurons typically express TNFα, and its expression intensifies in activated microglia and reactive astrocytes (Badoer, 2010). Fillit *et al.* disclosed meaningfully elevated levels of TNFα in the AD brains (Fillit et al., 1991). Animal studies confirm a link between excess TNFα levels in the brain and AD development (Janelsins et al., 2008).

We previously demonstrated that L-norvaline treatment led to a decline in IBA1 immunopositive cell’ density in the hippocampi of 3×Tg mice, which was associated with a shift from activated to resting ramified microglial phenotype. In the present study, we disclose a significant reduction in the hippocampal levels of TNFα mRNA expression following the treatment. It was convincingly demonstrated in primary cultures of mouse astrocytes that APP gene responds to the stimulation by TNFα (Lahiri et al., 2003). Remarkably, TNFα positively regulates APP transcription and translation via regulatory elements present in the APP promoter. Moreover, TNFα stimulates the expression levels of BACE1, which escalates the amyloidogenic APP processing (Zhao et al., 2011). Our findings confirm the direct connection between TNFα and APP but not BACE-1 levels in the 3×Tg mice.

There is a solidly grounded hypothesis that activated microglia cause synaptic and wiring dysfunction by pruning synaptic connections (Hong et al., 2016). Consequently, it was proposed to target the microglia-synapse pathways to prevent early AD symptoms (Xie et al., 2017). In the previous study, we reported a significant increase in dendritic spine density in the hippocampi of the L-norvaline treated mice. Here, we evidence an escalation of the PSD-95 protein levels in the hippocampi of the experimental group, which reflects the increase of spine density.

Accumulating empirical evidence suggests a role of mechanistic target of rapamycin (mTOR) in the regulation of immune responses (Powell et al., 2012) and major pathological processes of AD (Wang et al., 2014). Rapamycin improves memory and effectively reduces amyloid and tau pathologies in the 3×Tg mice (Caccamo et al., 2010). Accordingly, mTOR-signaling inhibition is a novel therapeutic target for AD (Tramutola et al., 2017). Recently, it was hypothesized and confirmed that mTOR affects TNF-mediated pro-inflammatory processes. Karonitsch *et al.* (2018) demonstrated in fibroblast culture that TNF activates the mTOR pathway, which, in turn, modulates the gene expression response to TNF (Karonitsch et al., 2018).

L-norvaline possesses anti-inflammatory properties, which have been attributed to its potency of inhibiting ribosomal protein S6 kinase beta-1 (S6K1). This kinase is a downstream target of mTOR signaling (Ming et al., 2009). Previously we reported a significant reduction (by 53%) of RAC-alpha protein-serine/threonine kinase (Akt1) levels in the L-norvaline treated mice (Polis et al., 2018). AKT1 is a key modulator of the AKT-mTOR-S6K1 signaling pathway, that regulates various biological processes including metabolism, proliferation, and cell survival (Memmott and Dennis, 2009). TNF was shown to induce the activation of AKT and S6K1 (Karonitsch et al., 2018). Therefore, we speculate that the reduction in the levels of AKT1 protein following the L-norvaline treatment is TNFα-mediated (Fig. 8). Of note, chronic rapamycin treatment does not modify the levels of TNFα in the brains of 3×Tg mice (Majumder et al., 2011). Thus, the observed reduction in the TNFα gene expression levels following the treatment cannot be explained by L-norvaline modulation of the AKT-mTOR-S6K1 signaling pathway. Future investigations should address the elucidation of this phenomenon.

**Figure 8.**
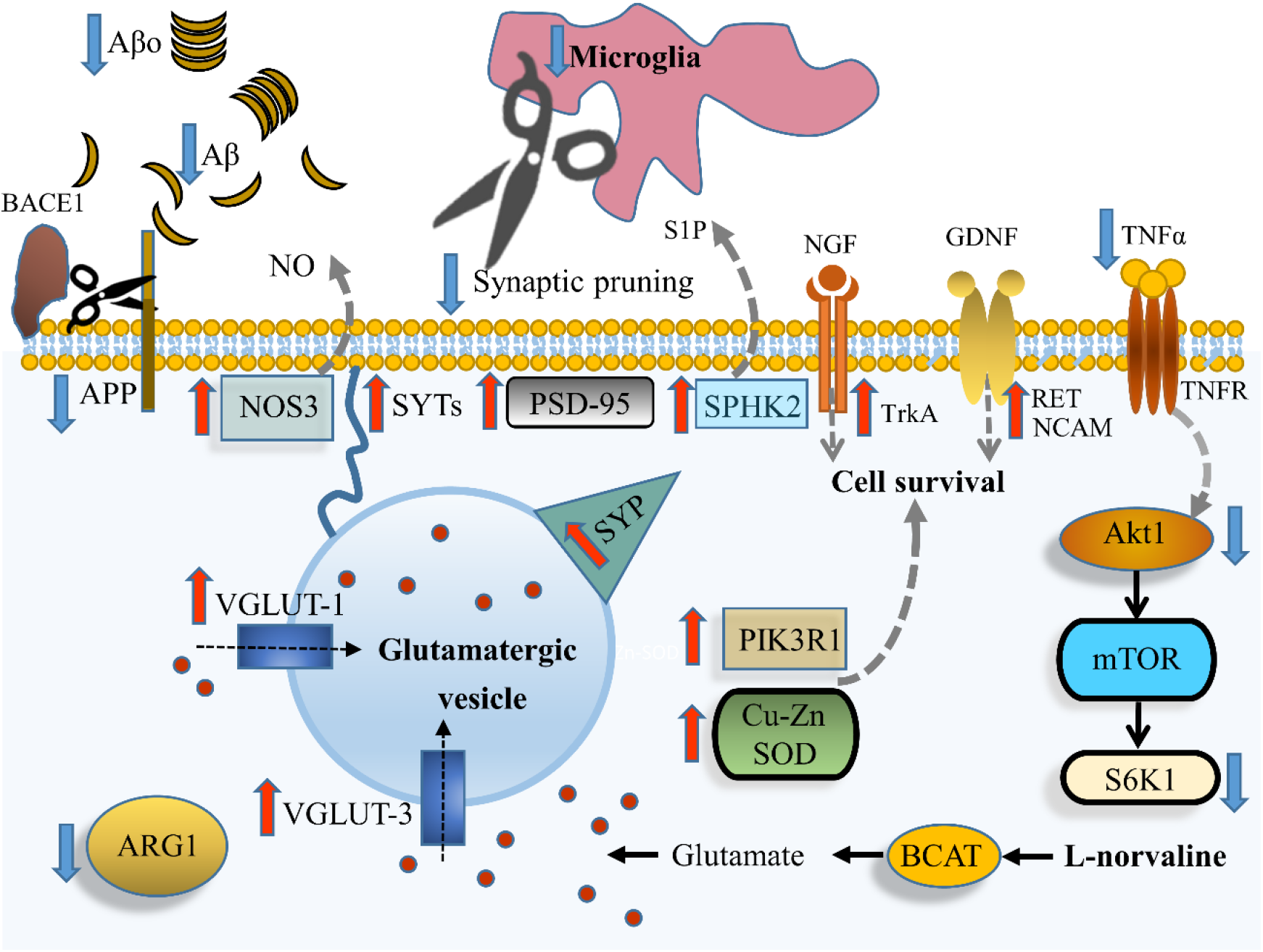
A proposed model for the metabolic effects of L-norvaline in the 3xTg mouse brain. The treatment leads to reduced quantities of Aβ fibrils and prefibrillar oligomers (Aβo), which is caused by a decline in the levels of APP but not BACE-1. L-norvaline treatment reduces microglial density and shifts their phenotype from activated to resting, which, in turn, decreases the levels of TNFα, reduces synaptic pruning and increases spine density. L-norvaline is a substrate of branched-chain amino acid aminotransferase (BCAT), which produces glutamate as the main product. The overproduction of glutamate leads to an increase in the expression levels of vesicular glutamate transporter 1 and 3 (VGLUT-1 and VGLUT-3, respectively), synaptophysin (SYP), synaptotagmins (SYTs). TNFα activates the mTOR pathway. L-norvaline also reduces the levels of Akt1 and inhibits S6K1. NGF binding toTrkA (which shows increased levels of expression in the L-norvaline treated mice) causes the phosphorylation of TrkA and activation of multiple preventing apoptosis signaling pathways. GDNF binds to a multi-component receptor (GDNFRα1 and RET), which induces cell survival mechanisms. The levels of RET are escalated in the L-norvaline treated mice. Additionally, NCAM (which is elevated by the treatment as well) serves as a second signaling receptor for GDNF. L-norvaline increases the levels of sphingosine kinase 2 (SPHK2), which leads to an escalation of the neuroprotective factor Sphingosine 1-phosphate (S1P) levels. Finally, L-norvaline increases the levels of superoxide dismutase [Cu-Zn] (SOD), and phosphatidylinositol 3-kinase regulatory subunit alpha (PIK3R1). Red arrows represent elevated levels, and blue arrows designate reduced levels compared to the vehicle-treated controls.

Our bioinformatics analysis revealed 41 critical biological processes involved in cell survival and immune response and significantly activated by L-norvaline treatment. The most significant pathways in the list are the Neuregulin (NRG) Signaling Pathway, Synaptic Long-Term Depression, ERK/MAPK Signaling, PDGF signaling, and Oncostatin M Signaling, which are essential for neuroprotection and their activation is particularly beneficial for AD patients.

L-norvaline is a potent non-competitive arginase inhibitor (Polis and Samson, 2018). Contemporary studies have identified arginase function in the brain and associated this enzyme with the development of neurodegenerative diseases (Patassini et al., 2015). Moreover, upregulation of arginase contributes to endothelial dysfunction, atherosclerosis, and diabetes. Therefore, regulation of arginase activity is an emerging universal approach for AD and other metabolic disorders treatment.

Previous studies established the presence of both ARG1 and ARG2 in the brain (Peters et al., 2013) (Polis et al., 2018). Moreover, arginase levels were shown to be increased in the areas with pronounced Aβ deposition (Kan et al., 2015), (Polis et al., 2018). Of note, the primary metabolic function of arginases in mammals is the removal of excess ammonia via the urea cycle. This is the central role of ARG1 in the hepatic tissue (Stewart and Caron, 1977). ARG2 is a kidney-type arginase, which is ubiquitously expressed at a low level within the mitochondria of various organs (Lange et al., 2004).

Arginase inhibition with a potent reversible inhibitor of liver arginase N(omega)-hydroxy-nor-L-arginine (nor-NOHA) (Ki=0.5 μM) ameliorates hepatic metabolism in obese mice (Moon et al., 2014). Of note, in the forementioned study, mice were orally gavaged with a relatively high dose of the arginase inhibitor (40 mg/kg per day) for five weeks. The authors conclude that arginase inhibition exerts protective properties against hepatic lipid abnormalities induced by obesity and address arginase inhibition as a potential therapeutic approach for obesity and its metabolic complications. Another study utilizing a several-fold higher (400–800 mg/kg) dose of nor-NOHA in rat demonstrated that arginase inhibition does not lead to changes in serum creatinine, alanine aminotransferase, alanine aminotransferase, or body weight in the animals. Moreover, histological analysis of liver, heart, lung, kidney, spleen, and pancreas did not reveal any structural abnormalities (Reid et al., 2007). Furthermore, even considerable reduction of the liver ARG1 activity (by 35%) does not affect general metabolic profile data in the rat (Sabbatini et al., 2003).

In this study, we aimed to investigate the influence of L-norvaline upon ARG1 and ARG2 levels in the primary organs of their expression. A set of the histochemical investigations revealed a significant reduction of ARG1 immunopositivity in the liver and ARG2 immunopositivity in the kidney following the treatment. Worth mentioning that ARG2 deficiency extends the lifespan of mice (Xiong et al., 2017). Moreover, targeting ARG2 protects mice from high-fat-diet-induced hepatic steatosis through suppression of macrophage inflammation (Liu et al., 2016). Interestingly enough, the release of TNFα from bone marrow-derived macrophages of Arg2^−/−^ mice is decreased as compared to the WT animals. Therefore, strong inhibition of ARG2 levels by L-norvaline might be beneficial for the experimental mice.

Previously we have proved that L-norvaline treatment does not lead to a significant drop in weight or detectable changes in the behavior of the WT mice (Polis et al., 2018). Moreover, the treatment does not affect the levels of ARG1 and ARG2 expression in their brains. Nevertheless, L-norvaline reduces amyloid-beta-driven arginase immunopositivity in the aged 3×Tg animals. For that reason, we suggest that arginase inhibition with L-norvaline provides a fine-tuning of the enzyme activity, primarily in the organs where some pathological stimuli upregulate it.

In conclusion, we emphasize that arginase inhibition, in general, shows wide-ranging therapeutic potential for the treatment of various pathologies. In this context, we underline multifaceted modes of L-norvaline activity, which interfere with several critical aspects of AD pathogenesis. Therefore, the substance represents a promising neuroprotective agent that deserves to be clinically investigated.

## Supporting information

## Acknowledgments

This research was supported by a Marie Curie CIG Grant 322113, a Leir Foundation Grant, a Ginzburg Family Foundation Grant, and a Katz Foundation Grant to AOS. We gratefully acknowledge Dr. Zohar Gavish for his help with immunohistochemistry and Dr. Tali Shalit for her help with bioinformatics analysis.

## Author Contributions

Baruh Polis and Abraham Samson designed the experiments. Baruh Polis performed the experiments and analyzed the data. Kolluru Devi Dutt Srikanth performed western blotting. Vyacheslav Gurevich performed RT-PCR. Hava Gil-Henn advised and supervised the experiments. Baruh Polis wrote the manuscript, and Abraham Samson and Hava Gil-Henn edited the manuscript.

