## Supplementary material for "L-Norvaline, a New Therapeutic Agent against Alzheimer’s disease"

| No | Target Name with Alias | Full Target Protein Name | %CFC | pvalue |
|---|---|---|---|---|
| **1** | SCNN1B | Amiloride-sensitive sodium channel subunit beta | 220 | 0.018 |
| **2** | Src | Src proto-oncogene-encoded protein-tyrosine kinase | 94 | 0.003 |
| **3** | MYPT1 | Protein phosphatase 1 regulatory subunit 12A | 94 | 0.009 |
| **4** | PP2A B | Protein-serine phosphatase 2A - B regulatory subunit - B56 alpha isoform | 92 | 0.032 |
| **5** | NPM1 (B23) | Nucleophosmin | 87 | 0.007 |
| **6** | MEK1 (MAP2K1; MKK1) | MAPK/ERK dual-specificity kinase 1 | 86 | 0.015 |
| **7** | PKCb (PRKCB1) | Protein-serine kinase C beta 1 | 85 | 0.017 |
| **8** | PLCG1 | 1-phosphatidylinositol 4,5-bisphosphate phosphodiesterase gamma-1 | 84 | 0.030 |
| **9** | Bcr | Breakpoint cluster region protein | 80 | 0.043 |
| **10** | Ros (ROS1) | Orosomucoid 1 receptor-tyrosine kinase | 78 | 0.003 |
| **11** | PHOCN (MOB4; mMOB1) | MOB-like protein phocein (Preimplantation protein 3) | 76 | 0.038 |
| **12** | STAT1a | Signal transducer and activator of transcription 1 alpha | 74 | 0.014 |
| **13** | ELK1 | ETS domain-containing protein Elk-1 | 72 | 0.032 |
| **14** | VGLUT3 | Vesicular glutamate transporter 3 | 71 | 0.012 |
| **15** | eNos (NOS3) | Nitric oxide synthase, endothelial | 68 | 0.041 |
| **16** | Rad17 | Cell cycle checkpoint protein RAD17 | 67 | 0.003 |
| **17** | Ksr1 | Protein-serine kinase suppressor of Ras 1 | 59 | 0.036 |
| **18** | TrkA (NGFR; NTRK1) | Nerve growth factor (NGF) receptor-tyrosine kinase | 56 | 0.039 |
| **19** | SPHK2 | Sphingosine kinase 2 | 56 | 0.036 |
| **20** | PKG1a (PRKG1A) | cGMP-dependent protein kinase I-alpha | 54 | 0.003 |
| **21** | Cyclin E1 (CCNE1) | Cyclin E1 | 54 | 0.015 |
| **22** | Ret (c-Ret; GDNF receptor; Glial cell line-derived neurotropic factor receptor) | Proto-oncogene tyrosine-protein kinase receptor Ret | 53 | 0.040 |
| **23** | Tlk1 | Tousled-like protein-serine kinase 1 | 53 | 0.029 |
| **24** | Synaptophysin | Synaptophysin | 50 | 0.039 |
| **25** | CaMKK (CaMKK1) | Calcium-calmodulin-dependent protein kinase kinase 1 | 50 | 0.026 |
| **26** | Haspin | Protein-serine/threonine kinase haspin | 49 | 0.033 |

| | | | | |
|---|---|---|---|---|
| **27** | PKCa (PRKCA) | Protein-serine kinase C alpha | 49 | 0.013 |
| **28** | MyoD (MYOD1) | Myoblast determination protein 1 | 47 | 0.035 |
| **29** | MEK1 (MAP2K1; MKK1) | MAPK/ERK dual-specificity kinase 1 | 46 | 0.001 |
| **30** | Mnk2 (MKNK2) | MAP kinase-interacting serine-threonine kinase 2 | 45 | 0.040 |
| **31** | STAT5A | Signal transducer and activator of transcription 5A | 45 | 0.006 |
| **32** | NrCAM | Neuronal cell adhesion molecule | 43 | 0.003 |
| **33** | INPP5F (OCRL) | Inositol polyphosphate 5-phosphatase OCRL-1 | 43 | 0.022 |
| **34** | PKAR2B | cAMP-dependent protein-serine kinase regulatory type 2 subunit beta | 43 | 0.038 |
| **35** | MRCKb | Protein-serine/threonine kinase MRCK beta | 42 | 0.001 |
| **36** | Mos | Moloney sarcoma oncogene-encoded protein-serine kinase | 42 | 0.012 |
| **37** | TNFR1 (CD120a; TNFRSF1A; TNFAR) | Tumour necrosis factor receptor superfamily member 1A | 42 | 0.022 |
| **38** | ACTA1 (Alpha-actin) | Actin, alpha, beta and gamma | 41 | 0.009 |
| **39** | Paxillin 1 (PXN) | Paxillin 1 | 41 | 0.034 |
| **40** | TEC | Protein-tyrosine kinase Tec | 40 | 0.034 |
| **41** | Nrf2 (NFE2L2) | Nuclear factor erythroid 2-related factor 2 | 40 | 0.020 |
| **42** | Beclin 1 (BECN1; GT197) | Beclin-1 | 40 | 0.045 |
| **43** | PP2A/Bb (PPP2R2B) | Protein-serine phosphatase 2A - B regulatory subunit - beta isoform | 39 | 0.016 |
| **44** | CavBeta2 (CACNB2; CAB2) | Voltage-dependent L-type calcium channel subunit beta-2 | 38 | 0.038 |
| **45** | ACACA (ACC1; | Acetyl-CoA carboxylase 1 | 37 | 0.032 |

| | | | | |
|---|---|---|---|---|
| | ACCA) | | | |
| **46** | DAB1 | Disabled homologue 1 | 37 | 0.015 |
| **47** | ACTA1 (Alpha-actin) | Actin, alpha skeletal muscle | 35 | 0.049 |
| **48** | Raf1 (c-Raf) | RAF proto-oncogene serine/threonine-protein kinase | 34 | 0.030 |
| **49** | Src | Src proto-oncogene-encoded protein-tyrosine kinase | 34 | 0.006 |
| **50** | BLNK | B-cell linker protein | 33 | 0.045 |
| **51** | SYT10 | Synaptotagmin-10 | 33 | 0.010 |
| **52** | STAT5B | Signal transducer and activator of transcription 5B | 32 | 0.019 |
| **53** | OSR1 | Protein odd-skipped-related 1 | 32 | 0.034 |
| **54** | Kit | Mast/stem cell growth factor receptor Kit | 31 | 0.029 |
| **55** | COX4I1 | Cytochrome c oxidase subunit 4 isoform 1, mitochondrial | 31 | 0.024 |
| **56** | PKCb2 (PRKCB2) | Protein-serine kinase C beta 2 | 30 | 0.047 |
| **57** | PKCb (PRKCB1) | Protein-serine kinase C beta 1 | 29 | 0.041 |
| **58** | MOK | MAPK/MAK/MRK overlapping kinase | 29 | 0.040 |
| **59** | PKR1 (PRKR; EIF2AK2) | Double stranded RNA dependent protein-serine kinase | 29 | 0.039 |
| **60** | RAB5A (Rab5) | Ras-related protein Rab-5A | 29 | 0.018 |
| **61** | HSP105 (HSPH1, HSP110) | Heat shock 105 kDa protein | 29 | 0.032 |
| **62** | CaMK1a (CaMKI) | Calcium/calmodulin-dependent protein-serine kinase 1 alpha | 29 | 0.003 |
| **63** | MEK2 (MAP2K2; MKK2) | MAPK/ERK dual-specificity kinase 2 | 29 | 0.037 |
| **64** | EphA1 | Ephrin type-A receptor 1 protein-tyrosine kinase | 28 | 0.041 |
| **65** | CaMK1a (CaMKI) | Calcium/calmodulin-dependent protein-serine kinase 1 alpha | 28 | 0.043 |
| **66** | MKK3 (MAP2K3; MEK3) | MAPK/ERK dual-specificity kinase 3 beta isoform | 28 | 0.003 |
| **67** | PAK3 (PAKb) | p21-activated kinase 3 (beta) (Protein-serine/threonine kinase PAK3) | 27 | 0.046 |
| **68** | Cyclin D1 (CCND1) | Cyclin D1 | 25 | 0.031 |
| **69** | JAK3 | Janus protein-tyrosine kinase 3 | 25 | 0.030 |
| **70** | PP2A/Ca (PPP2CA)/'PP2A/Cb (PPP2CB) | Protein-serine phosphatase 2A - catalytic subunit - alpha and beta isoform | 24 | 0.022 |
| **71** | PP2B-B1/2 | Calcineurin subunit B type 2 | 24 | 0.005 |
| **72** | PIK3R1 (PI3K p85) | Phosphatidylinositol 3-kinase regulatory subunit alpha | 24 | 0.034 |

| | | | | |
|---|---|---|---|---|
| **73** | Synapsin 1 | Synapsin 1 isoform Ia | 24 | 0.009 |
| **74** | RSK4 (RPS6KA6) | Ribosomal S6 protein-serine kinase 4 (alpha 6) | 24 | 0.001 |
| **75** | ITGA4 (CD49D) | Integrin alpha 4 (VLA4) | 23 | 0.044 |
| **76** | KRAS (KRAS2) | GTPase KRas | 23 | 0.002 |
| **77** | AMPKa1 | 5'-AMP-activated protein kinase catalytic subunit alpha-1 | 22 | 0.035 |
| **78** | Pim2 | Protein-serine/threonine kinase pim-2 | 22 | 0.003 |
| **79** | DICER1 | Endoribonuclease Dicer | 21 | 0.028 |
| **80** | p38d MAPK (MAPK13) | Mitogen-activated protein-serine kinase p38 delta | 20 | 0.023 |
| **81** | VEGFR1 (Flt1) | Vascular endothelial growth factor receptor 1 | 20 | 0.026 |
| **82** | LAR (PTPRF) | Receptor-type tyrosine-protein phosphatase F | 20 | 0.002 |
| **83** | WNK2 (PRKWNK2) | Protein-serine/threonine kinase WNK2 | 20 | 0.026 |
| **84** | ATM | Ataxia telangiectasia mutated | 20 | 0.011 |
| **85** | SOD3 | Extracellular superoxide dismutase [Cu-Zn] | 19 | 0.041 |
| **86** | CDK10 | Cyclin-dependent protein-serine kinase 10 | 19 | 0.019 |
| **87** | TRRAP | Transformation/transcription domain-associated protein | 18 | 0.037 |
| **88** | PUMA (CHST9) | Carbohydrate sulfotransferase 9 | 18 | 0.048 |
| **89** | MRCKa | Protein-serine/threonine kinase MRCK alpha | 18 | 0.018 |
| **90** | GSK3 Beta (GSK3b) | Glycogen synthase kinase-3 beta | 18 | 0.017 |
| **91** | BCL2A1 | Bcl-2-related protein A1 | 17 | 0.049 |
| **92** | PP1/Ca (PPP1CA) | Protein-serine phosphatase 1 - catalytic subunit - | 17 | 0.009 |

| | | | | |
|---|---|---|---|---|
| | | alpha isoform | | |
| **93** | TLR4 (CD284) | Toll-like receptor 4 | 17 | 0.037 |
| **94** | JNK3 (SAPKb) | Jun N-terminus protein-serine kinase 3 (Stress-activated protein kinase-beta) | 16 | 0.036 |
| **95** | SLC38A1 (ATA1; NAT2) | Sodium-coupled neutral amino acid transporter 1 | 16 | 0.049 |
| **96** | SYT6 | Synaptotagmin-6 | 16 | 0.014 |
| **97** | ATG4B | Cysteine protease, Autophagy-related protein ATG4B | 15 | 0.032 |
| **98** | CDKL3 | Cyclin-dependent kinase-like 3 | 15 | 0.044 |
| **99** | VEGFR2 (KDR, Flk1) | Vascular endothelial growth factor receptor-tyrosine kinase 2 | 14 | 0.028 |
| **100** | Malin (NHLRC1; EPM2B) | E3 ubiquitin-protein ligase NHLRC1 | 14 | 0.033 |
| **101** | JNK1 (MAPK8; SAPK1) | Jun N-terminus protein-serine kinase (Stress-activated protein kinase) 1 | 14 | 0.000 |
| **102** | PKCm (PRKCM, PRKD1, PKD1) | Protein-serine kinase C mu (Protein kinase D) | 14 | 0.039 |
| **103** | MKK3 (MAP2K3; MEK3) | MAPK/ERK dual-specificity kinase 3 beta isoform | 14 | 0.037 |
| **104** | SRPK2 | Serine/arginine-rich protein-specific kinase 2 (SRSF protein kinase 2) | 13 | 0.007 |
| **105** | JAK1 | Janus protein-tyrosine kinase 1 | 13 | 0.013 |
| **106** | SYT10 | Synaptotagmin-12 | 12 | 0.032 |
| **107** | SGK1 | Protein-serine/threonine kinase Sgk1 | 11 | 0.028 |
| **108** | Caveolin 2 | Caveolin 2 | 11 | 0.038 |
| **109** | PP2A/Bb (PPP2R2B) | Protein-serine phosphatase 2A - B regulatory subunit - beta isoform | 10 | 0.010 |
| **110** | AurKA (Aurora A, AIK) | Aurora Kinase A (Protein-serine/threonine kinase 6) | 10 | 0.021 |
| **111** | CD45 | Receptor protein-tyrosine phosphatase CD45 | 9 | 0.042 |
| **112** | TAK1 (MAP3K7) | TGF-beta-activated protein-serine kinase 1 | 9 | 0.018 |
| **113** | PDK1 (PDHK1) | [Pyruvate dehydrogenase [lipoamide]] kinase isozyme 1, mitochondrial | 8 | 0.017 |
| **114** | ULK4 | Unc-51-like protein-serine/threonine kinase 4 | 3 | 0.005 |
| **115** | p38g MAPK (MAPK12, ERK6, SAPK3) | Mitogen-activated protein-serine kinase p38 gamma | -10 | 0.015 |
| **116** | CK1e (CSNK1E) | Casein protein-serine kinase 1 epsilon | -10 | 0.027 |
| **117** | SCN3B (NaVbeta3) | Sodium channel subunit beta-3 | -11 | 0.041 |

| | | | | |
|---|---|---|---|---|
| **118** | PTGES3 (p23) | Prostaglandin E synthase 3 | -13 | 0.005 |
| **119** | A-Raf | Protein-serine/threonine kinase A-Raf | -15 | 0.031 |
| **120** | Nek2 | NIMA (never-in-mitosis)-related protein-serine kinase 2 | -16 | 0.025 |
| **121** | Striatin | Striatin | -22 | 0.015 |
| **122** | HO1 (HO; HMOX1) | Heme oxygenase 1 | -24 | 0.005 |
| **123** | REEP1 | Receptor expression-enhancing protein 1 | -25 | 0.049 |
| **124** | Plk4 (SAK; STK18) | Polo-like protein-serine kinase 4 | -25 | 0.004 |
| **125** | DUSP7 | Dual specificity protein phosphatase-7 | -25 | 0.003 |
| **126** | HSP72 | Heat shock-related 70 kDa protein 2 | -27 | 0.024 |
| **127** | HSP90AB1 (HSP90; HSP84; HSP90B; HSPC2; HSPCB) | Heat shock 90 kDa protein beta | -35 | 0.045 |
| **128** | PP2A B (PPP2R5A; B56) | Protein-serine phosphatase 2A - B regulatory subunit - B56 alpha isoform | -36 | 0.008 |
| **129** | CDK5 | Cyclin-dependent protein-serine kinase 5 | -36 | 0.024 |
| **130** | PDI (P4hb; PDIA1; ERBA2L; PO4DB) | Protein disulfide-isomerase | -38 | 0.042 |
| **131** | JAK2 | Janus protein-tyrosine kinase 2 | -39 | 0.043 |
| **132** | FIH (HIF1; HIF1AN) | Hypoxia-inducible factor 1-alpha inhibitor | -45 | 0.017 |
| **133** | TRPC5 (TRP5) | Short transient receptor potential channel 5 | -47 | 0.029 |
| **134** | DUSP5 | Dual specificity protein phosphatase-5 | -49 | 0.040 |
| **135** | WNK3 (PRKWNK3) | Protein-serine/threonine kinase WNK3 | -51 | 0.045 |
| **136** | Akt1 (PKBa) | RAC-alpha protein-serine/threonine kinase | -53 | 0.038 |
| **137** | Grp75 (HspA9) | Stress-70 protein, mitochondrial | -59 | 0.008 |
